# Speech-driven Facial Animations Improve Speech-in-Noise Comprehension of Humans

**DOI:** 10.1101/2021.12.18.471222

**Authors:** Enrico Varano, Konstantinos Vougioukas, Pingchuan Ma, Stavros Petridis, Maja Pantic, Tobias Reichenbach

## Abstract

Understanding speech becomes a demanding task when the environment is noisy. Comprehension of speech in noise can be substantially improved by looking at the speaker’s face, and this audiovisual benefit is even more pronounced in people with hearing impairment. Recent advances in AI have allowed to synthesize photorealistic talking faces from a speech recording and a still image of a person’s face in an end-to-end manner. However, it has remained unknown whether such facial animations improve speech-in-noise comprehension. Here we consider facial animations produced by a recently introduced generative adversarial network (GAN), and show that humans cannot distinguish between the synthesized and the natural videos. Importantly, we then show that the end-to-end synthesized videos significantly aid humans in understanding speech in noise, although the natural facial motions yield a yet higher audiovisual benefit. We further find that an audiovisual speech recognizer benefits from the synthesized facial animations as well. Our results suggest that synthesizing facial motions from speech can be used to aid speech comprehension in difficult listening environments.

## 1 Introduction

Real-world listening environments are often noisy: many people talk simultaneously in a busy pub or restaurant, background music plays frequently, and traffic noise is omnipresent in cities. Seeing a speaker’s face makes it considerably easier to understand them (Sumby and Pollack 1954, Ross et al. 2007), and this is particularly true for people with hearing impairments (Puschmann et al. 2019) or who are listening in background noise. This phenomenon, termed inverse effectiveness, is characterized by a more pronounced audiovisual comprehension gain in challenging hearing conditions (Crosse et al. 2016, Stevenson and James 2009, Meredith and Stein 1986).

This audiovisual (AV) gain is linked to the temporal and categorical cues carried by the movement of the head, lips, teeth, and tongue of the speaker (Chandrasekaran et al. 2009, O’Sullivan et al. 2017, Munhall et al. 2004) and likely emerges from multi-stage, hierarchical predictive coupling and feedback between the visual and the auditory cortices (Peelle and Sommers 2015, Hickok and Poeppel 2007, Kayser et al. 2012, Schroeder et al. 2008, Kayser et al. 2007, O’Sullivan et al. 2021, Crosse et al. 2016).

However, the visual component of audiovisual speech is often not available, such as when talking on the phone or to someone wearing a mask, when listening to the radio or when watching video content where the audio narrates non-speech video content. A system that automatically synthesizes talking faces from speech and presents them to a listener could potentially aid comprehension both for normal hearing people and those living with hearing loss in such situations.

Early efforts to synthesize talking faces from speech were based on pre-recorded kinematic and parametrized models (Kuratate et al. 1998, Cohen and Massaro 1990). These early models yielded animations capable of augmenting speech comprehension in background noise (Munhall et al. 2004, Le Goff et al. 1997, Massaro and Cohen 1990) but required the previous or simultaneous recording of a human speaker wearing facial markers or electromyographic electrodes (Bailly et al. 2003).

Later works proposed a modular framework for pre-trained text-to-AV-speech synthesizers (MASSY) which included both animated and photorealistic face generation sub-modules (Fagel and Sendlmeier 2003, Fagel 2004). Talking heads synthesized with such models increased comprehension performance as much as their natural counterparts in consonant-recognition paradigms but word and sentence identification was about twice as high for the natural videos (Aller and Meister 2016, Lidestam and Beskow 2006).

Synface, a project dedicated to synthesizing talking faces for enhancing speech comprehension, also utilized phonetic analysis of speech and showed that stimuli generated in such a way can improve speech comprehension in people with hearing impairments as well as in healthy volunteers listening in background noise (Agelfors et al 2006, Beskow et al 2002).

Recent advances in speech-driven animation methods have made it possible to produce photorealistic talking heads with synchronized lip movements using only a still image and an audio clip. State-of-the-art solutions are trained in an end-to-end manner using self-supervision and do not require intermediate linguistic features such as phonemes, or visual features such as facial landmarks and visemes. Most are based on generative adversarial networks (GANs) and can produce high quality visual signals that can even reflect the speaker’s emotion (Chung et al. 2017, Chen et al. 2019, Vougioukas et al. 2020).

Employing such facial animations to improve speech-in-noise comprehension would represent a significant step forward in the development of audiovisual hearing aids. However, it has not yet been investigated whether such end-to-end synthetic facial animations can aid a listener to better understand speech in noisy backgrounds. In this study we set out to investigate this issue.

## 2 Material and Methods

To investigate the impact of different types of AV speech on speech-in-noise comprehension in humans, we first synthesized realistic facial animations from speech. We then assessed how these facial animations benefitted humans in understanding speech in noise, compared to no visual signal and to the actual video of a speaker. We finally compared the human level of AV speech comprehension to that of an AV automatic speech recognizer.

### 2.1 Audiovisual material

We employed sentences from the GRID corpus, which consists of 33 speakers each uttering 1,000 three-second-long sentences (Cooke et al. 2006). The videos in the GRID corpus are recorded at 25 frames per second, and the speech signals are sampled at 50 kHz. Four speakers, of which two were female, were selected for their lack of a strong accent (speakers 12, 19, 24 and 29).

Sentences of the GRID corpus are semantically unpredictable but meaningful commands composed of six words taken from a limited dictionary (Table 1). As intended for this corpus, participants were only scored on the color, letter, and digit in each sentence (i.e., the keywords marked with an asterisk in Table 1), with the remaining words acting as contextual cues.

**Table 1:** Structure of GRID corpus sentences. The keywords on which participants were scored are indicated by an asterisk (*).

| Command | Color* | Preposition | Letter* | Digit* | Adverb |
| --- | --- | --- | --- | --- | --- |
| Bin | blue | at | a-z | 0-9 | again |
| Lay | green | by | *except* |  | now |
| Place | red | in | w |  | soon |
| Set | white | with |  |  | please |

#### 2.1.1 Audio

The audio files of the chosen speakers were down-sampled to 48 kHz using FFMPEG to match the sampling frequency of the available speech-shaped noise (SSN) files. The latter, also known as speech-weighted noise, was generated from the spectral properties of multiple concatenated clean speech files from different speech corpora and audiobooks by randomizing the phase of all spectral components before extracting the real part of the inverse Fourier transform.

The root mean square amplitudes of both the voiced part of the GRID sentence and the SSN were then measured. The two signals were scaled and combined such that the signal-to-noise ratio (SNR) was −8.82dB. This value was found during pilot testing to reduce comprehension of normal-hearing participants to 50%.

#### 2.1.2 Synthesized Video

We used the GAN^1^ model proposed by Vougioukas et al. (2020) to generate talking head videos from single still images and speech signals at 25 frames per second (Figure 1). The GAN is trained using multiple discriminators to enforce different aspects of realism on the generated videos, including a synchronization discriminator for audiovisual synchrony. The offset between the audio and the visual component in the synthesized videos is below 1 frame (below 40 ms, Table 6, Vougioukas et al. 2020). This method is also capable of generating videos that exhibit spontaneous facial expressions such as blinks, which contribute to the realism of the sequences.

**Figure 1:**
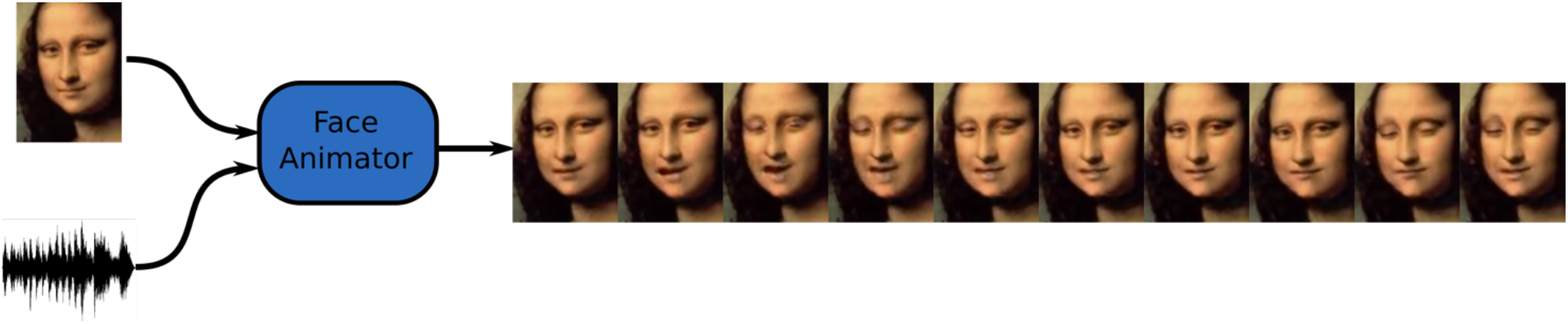
Schematic of the generation of the facial animation. A GAN synthesizes a video of a talking face from a still image of the speaker and a speech signal. N.B.: we include such images for illustration purposes only: the real/generated GRID images were shown to participants in the study.

The LipNet pretrained automated lipreading model, which obtains a word error rate (WER) of 21.76% on the natural images, achieves a WER performance of 23.1% when evaluated on synthetic videos from unseen subjects of the GRID dataset, indicating that the produced movements correspond to the correct words (Vougioukas et al. 2020, Assael et al. 2016).

#### 2.1.3 Natural Video

For direct comparability with the synthesized videos, the natural videos presented to the volunteers were formatted in the same way as the natural videos used to train the GAN. The faces in the high-resolution GRID videos were aligned to the canonical face, cropped, and downscaled to a resolution of 96×128 pixels using FFMPEG. The points at the edges of the eyes and tip of the nose were used for the alignment of the face. The process used to obtain videos focused on the face is outlined in Figure 2.

**Figure 2:**
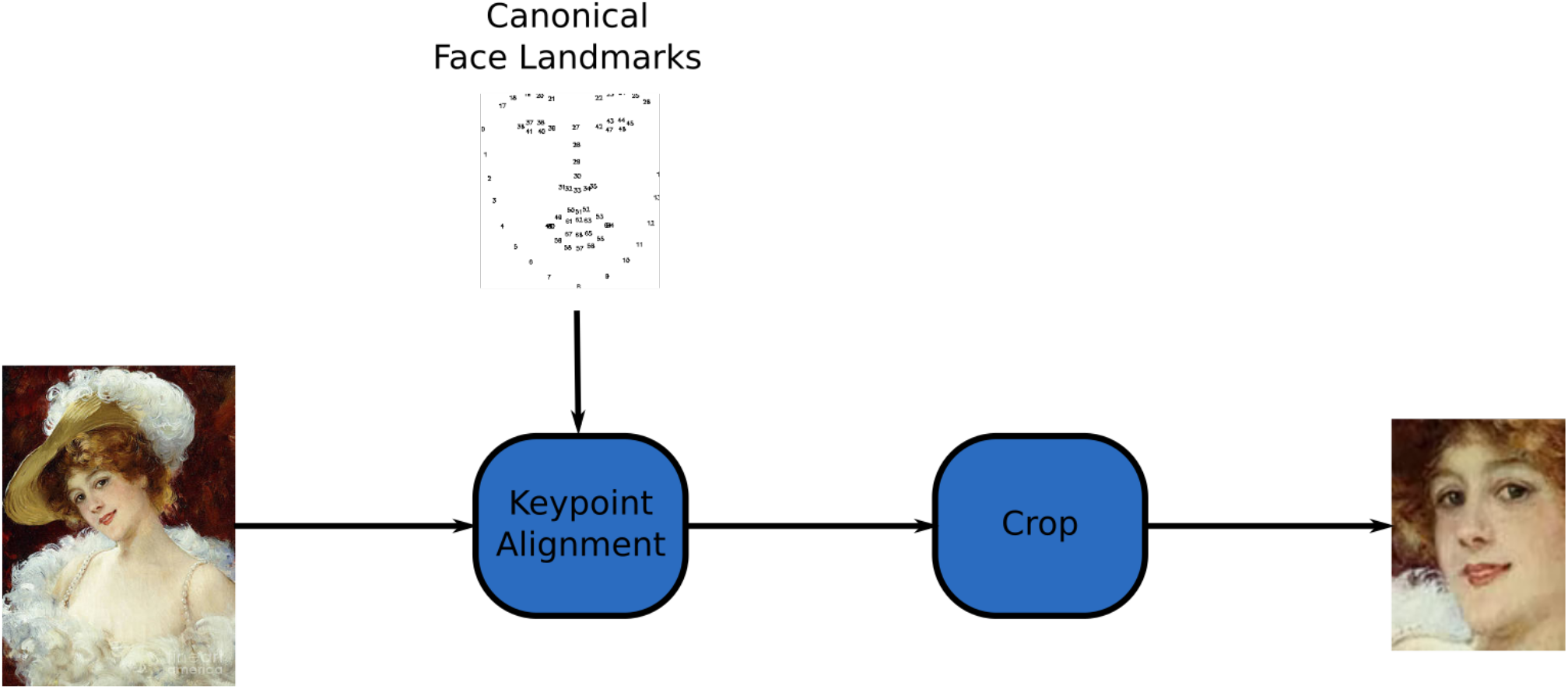
Video preprocessing pipeline used to obtain cropped videos of the speaker’s face. N.B.: we include such images for illustration purposes only: the real/generated GRID images were shown to participants in the study.

### 2.2 Turing Realism Test

The realism of the synthesized videos was assessed through an online Turing test. Users were shown 24 randomly selected videos from the GRID, TIMIT (Garofolo et al. 1993) and CREMA (Cao et al. 2014) datasets, half of which were synthesized, and were asked to label them as real or fake in a two-alternative forced choice (2AFC) procedure. The experiment was performed by 50 students and staff members from Imperial College London before the Turing test was made available online^2^. The results from the first 750 respondents were reported in Vougioukas et al. (2020) and we present updated results from 1,217 participants. Figure 3 shows a side-by-side comparison between a fake and generated video.

**Figure 3:**
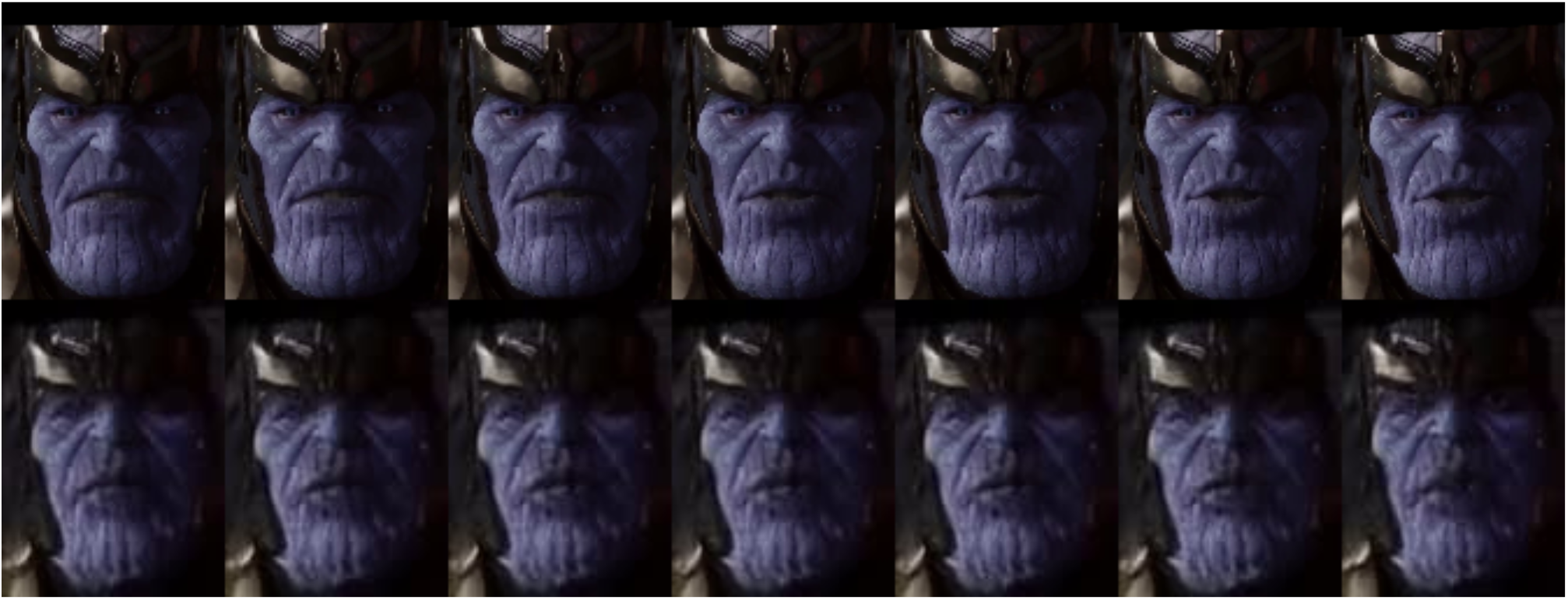
An example of frames from a generated video (top row) shown alongside the corresponding real video (bottom row). NB: we include such images for illustration purposes only: the real/generated GRID images were shown to participants in the study.

An unstructured assessment of the videos’ realism was also performed on the 18 participants of the speech-comprehension experiment (see below). Following the speech comprehension task, the subjects were asked to comment on anything interesting or strange they had noticed in the videos during the experiment. Their verbal responses were recorded anonymously.

### 2.3 Assessment of speech-in-noise comprehension

#### 2.3.1 Participants

Eighteen native English speakers, eleven of them female, with self-reported normal hearing and normal or corrected-to-normal vision participated in the experiment. The participants were between 18 and 36 years of age, with a mean age of 23 years. All participants were right-handed and had no history of mental health problems, severe head injury or neurological disorders. Before starting the experiment, participants gave informed consent. The experimental protocol was approved by the Imperial College Research Ethics Committee.

#### 2.3.2 Stimuli presentation

We considered three types of AV stimuli. All three types had speech in a constant level of background noise, that is, with the same SNR. The type of the video, however, varied between the three types of AV signals. During one type of stimulation, subjects heard noisy speech while the monitor remained blank (“audio-only”). In another type, we presented subjects with noisy speech together with the synthesized facial animations (“synthetic AV”). Finally, subjects were also presented with the speech signals while watching the genuine corresponding videos of the talking faces (“natural AV”).

The experiment consisted of six rounds of three blocks, where each block corresponded to one of the AV conditions. Six sentences were presented in each block. The order in which the three conditions were presented was randomized within rounds and across rounds. Each sentence was chosen randomly from a pool of all 1,000 sentences from each of the four speakers, and the order of speakers was randomized.

Each subject therefore listened to 36 sentences for each of the three AV types. The participants took a brief rest for one minute after every round.

#### 2.3.3 Data and analysis

Between each sentence, the participants were asked to select the keywords they had heard, from a list on the screen. The list allowed participants to select all possible GRID sentence combinations while non-keyword terms were pre-selected and displayed for them in each trial. The selection of the keywords by the participants on the monitor allowed to compute their comprehension score automatically.

The data was therefore collected and analyzed in a double-blind fashion: neither the experimenter nor the participant knew which type of video or what specific sentence was presented. Importantly, the participants were not informed of the synthesized nature of part of the videos.

The scoring was expressed as the percentage of keywords correctly identified in each trial. The scores for each type of AV signal were extracted by averaging across trials and rounds for each participant. The responses for each keyword were also recorded, paired with the corresponding presented keyword.

#### 2.3.4 Hardware and software

The experiment took place in a an acoustically and electrically insulated room (IAC Acoustics, UK). A computer running Windows 10 placed outside the room controlled the audiovisual presentation and data acquisition. The audio component of the stimulus was delivered diotically at a level of 70 dB(A) SPL using ER-3C insert earphones (Etymotic, USA) through a high-performance sound card (Xonar Essence STX, Asus, USA). The sound level was calibrated with a Type 4157 ear simulator (Brüel&Kjær, DK). The videos were delivered through a fast 144 Hz, 24-inch monitor (24GM79G, LG, South Korea) set at a refresh rate of 119.88 Hz. The monitor was mounted at a distance of one meter from the participant. The videos were played in full screen such that the dimensions of the talking heads appeared life-sized.

To ensure that the audio and video components of the stimuli were presented in synchrony, the audiovisual latency of the presentation system was characterized. A photodiode (Photo Sensor, BrainProducts, Germany) attached to the display and an acoustic adaptor (StimTrak, BrainProducts, Germany) attached to the audio cable that was connected to the ear phones were employed to record the output of a prototypical audiovisual stimulus. The latency difference between the two stimuli modalities was found to be below 8 ms.

### 2.4 Audiovisual automated speech recognition

The same 36 sentences that were randomly selected and presented to each participant for each condition were also analyzed with an audiovisual speech recognizer (AVSR). We fine-tuned the pre-trained model from Ma et al. (2021) for ten epochs on the 29 GRID speakers which were not used in the behavioral study. The AVSR employed ResNets to extract features directly from the mouth region coupled with a hybrid connectionist temporal classification (CTC) objective/attention architecture. The output of the model was then analyzed in the same way as the human data.

## 3 Results

To assess the realism of our facial animations, we first investigated whether humans could discriminate between the synthesized videos and the natural ones. In a large online Turing test on 1,217 subjects, we found that the median of the correct responses was exactly at the chance level of 50% (Figure 4), as was the result on a more controlled Turing test performed on 50 subjects. Moreover, the 18 participants of the speech-in-noise comprehension experiment were not told of the nature of half of the videos, and none reported finding anything unusual regarding the videos in a questionnaire completed following the experiment. To the average human observer, the synthesized videos were thus indistinguishable from the natural ones.

**Figure 4:**
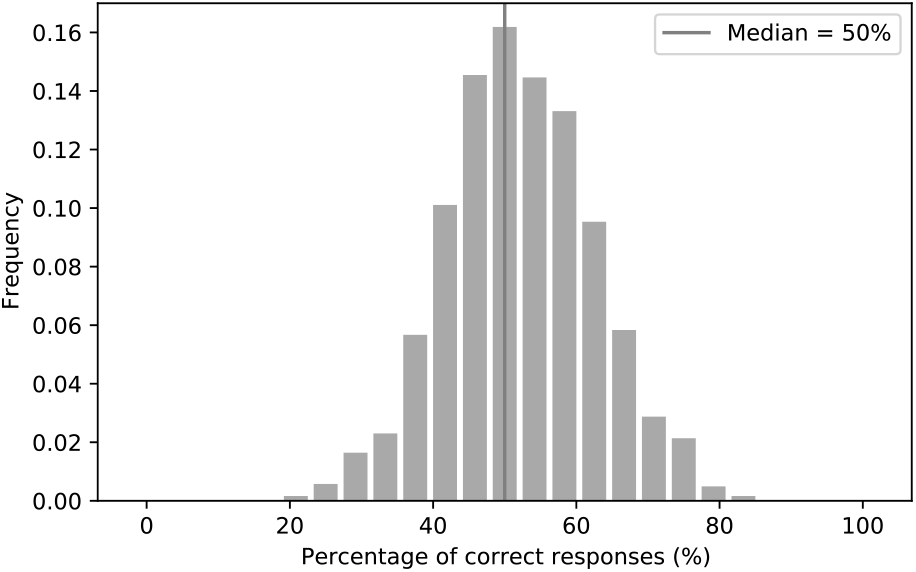
Histogram of the percentage of correct responses in the Turing test on discriminating between the synthetic and the natural videos. The median was exactly at the chance level of 50%.

We then proceeded to assess the potential benefits of the synthesized talking faces on speech-in-noise comprehension. We found that both the synthesized and the natural videos significantly improved comprehension in our participants when compared to the audio signal alone (Figure 5).

**Figure 5:**
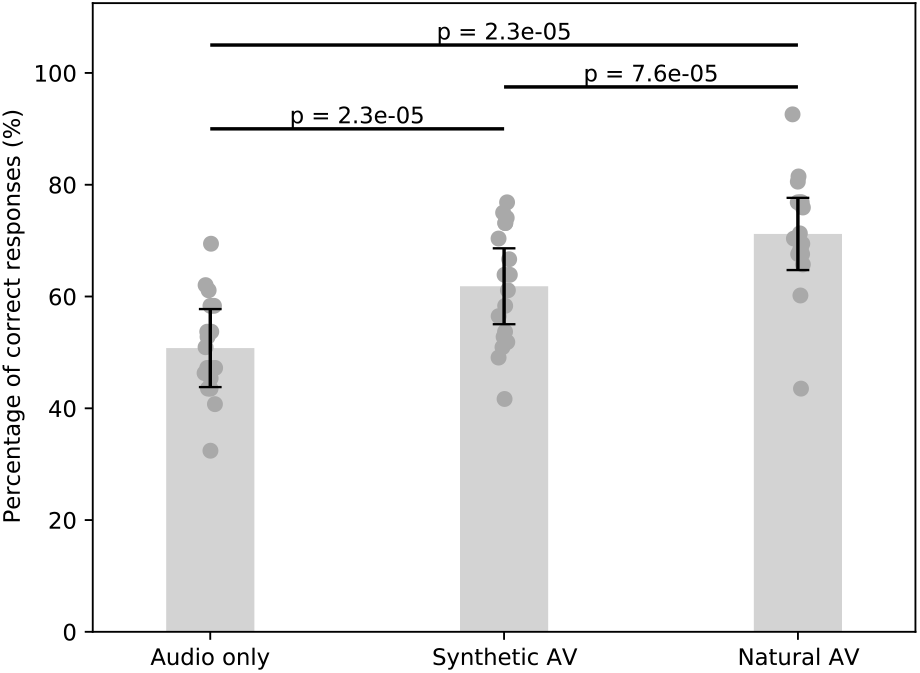
Speech comprehension in noise without a visual signal, with the synthesized facial animations, as well as with the natural AV signals. Error bars represent the standard error of the mean, and the dark grey points show the average score per subject.

The comprehension for the audio-only type was 50.8% ± 7% (mean and standard error of the mean). The synthesized and natural videos improved speech comprehension to 61.8% ± 7% and 71.2% ± 6% respectively. The relative improvement between the audio-only and synthetic AV signals was about 22% (p = 2.3×10^−5^, w=1, two-sided Wilcoxon signed-rank test for dependent data with Benjamini–Hochberg FDR correction). The relative improvement of the natural AV signals as compared to the audio signal alone was about twice as large, about 40% (p = 2.3×10^−5^, w=0). The relative difference between the synthetic AV signals and the natural ones was statistically significant as well, at 15% (p = 7.6×10^−5^, w=5).

We further analyzed the differences between the AV gain in speech comprehension provided by the synthetic and the natural AV signals. In particular, we computed confusion matrices between the different key words of the sentences that the volunteers were asked to understand. The confusion matrices were normalized such that, for each particular keyword, the probability to select any other keyword was one. We then subtracted answer-response pair frequency of the confusion matrix of the synthesized AV signals from that of the natural AV signals (Figure 6A). As indicated by the presence of mostly positive differences on the leading diagonal of the resulting matrix, the natural videos outperformed the synthesized videos in terms of providing categorically unequivocable cues. The differences in the remaining sectors of the matrix shed some light into the reason the natural videos performed better. For example, matrix elements highlighted by the green rectangle in Figure 6A demonstrate that the synthesized videos encouraged participants to mistakenly select the letter ‘*a*’ when presented with the keywords ‘*o*’ and ‘*n*’. Similarly, the yellow arrows highlight that participants were more likely to mistake the letter ‘*t*’ for the letter ‘*g*’ and the digit ‘*two*’ for the digit ‘*seven*’ when presented with synthesized videos relatively to the natural videos.

**Figure 6:**
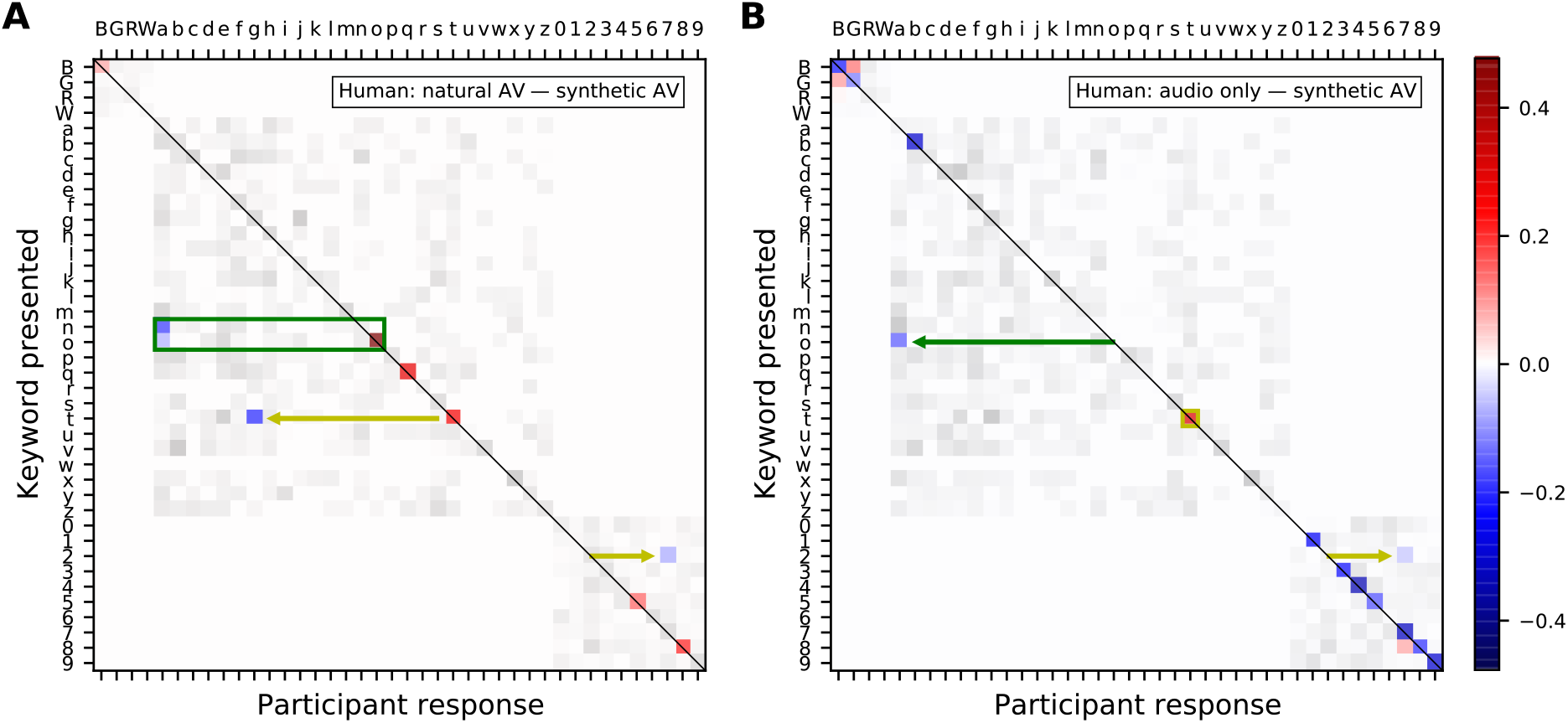
Confusion matrices showing the difference in frequency of presented keyword and participant response pairs between A) the natural audiovisual condition and the synthesized audiovisual condition and B) the audio-only condition and the synthesized audiovisual condition, in humans. The color keywords are represented by their first letter in uppercase, those corresponding to a letter are shown in lowercase, and the number keywords by their digit. The magnitude of statistically significant pairs is highlighted in color (bootstrapped permutation test, alpha=0.05, FDR corrected). A blue color indicates that subjects were significantly more likely to select a particular keyword when being presented with the synthesized visual signal as opposed to A) the natural video and B) no video.

We also subtracted the answer-response pair frequency of the confusion matrix of the synthetic AV signals from that obtained from audio-only signals (Figure 6B). The mostly negative differences on the leading diagonal of the resulting matrix show that the synthetic videos improved the subjects’ ability to discriminate between keywords compared to the audio-only condition. The green arrow in Figure 6B highlights one exception: the synthetic videos encouraged participants to mistakenly select the keyword ‘*a*’ when presented with ‘o’, congruently with the results shown in panel A. The yellow annotations indicates that the confusion of the keywords ‘t’ and ‘two’ also persists.

Nonetheless, the synthesized videos were found to disambiguate the keyword ‘*b*’, notable for being hard to distinguish from other consonants pronounced in combination with the phoneme /i:/ such as the keywords ‘*g*’ and ‘*d*’.

We then determined whether an AVSR could benefit from the synthetic facial animations as well. We found that the scores of the AVSR improved by about 13% for the synthetic AV material as compared to the audio-only signals (Figure 7). However, this improvement was significantly lower than the corresponding improvement of 22% in human speech-in-noise comprehension (p = 0.007, t=3.04, two-sided single-value t-test). Also, the natural AV signals improved the scores of the AVSR by 40% when compared to the audio signal alone, which was comparable to our result on the gain in human speech comprehension.

**Figure 7:**
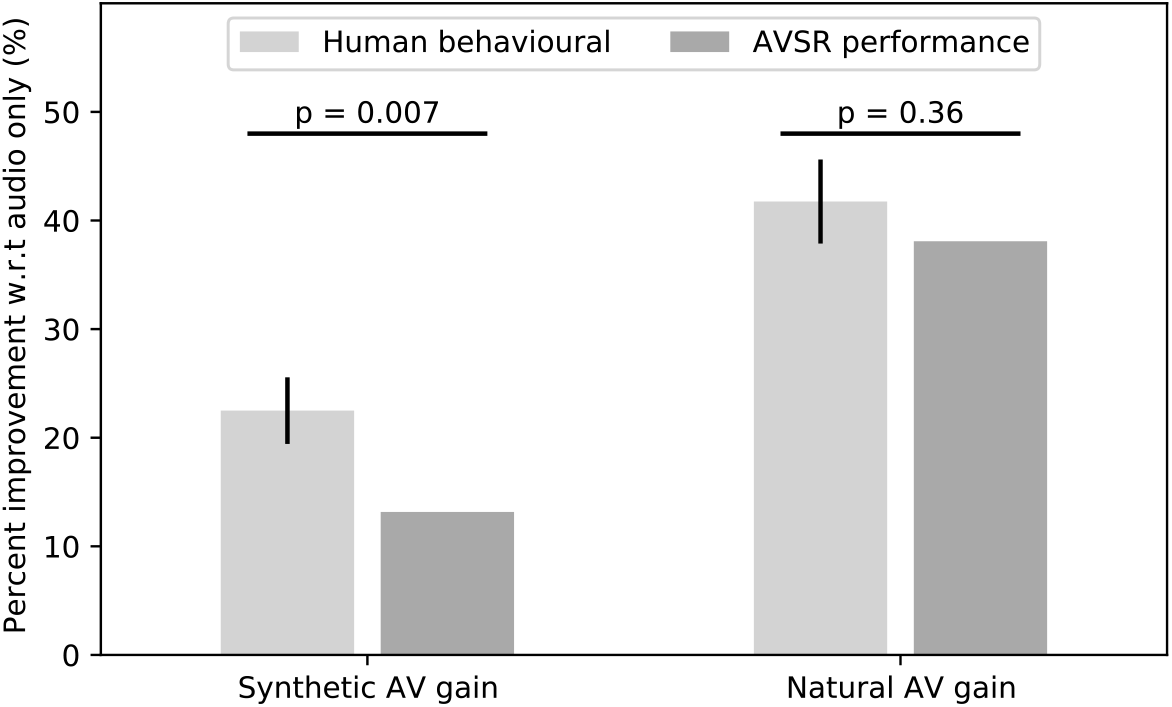
The scores of an AVSR were improved by the synthetic videos, although the gain was less than that experienced by humans. The natural AV signals led to a higher gain, similar to that of our human volunteers.

We also analyzed the confusion matrices for the AVSR data (Figure 8), which were calculated in the same way as those for the human behavioral data. The natural videos outperformed the synthetic videos across most keywords, in particular allowing the AVSR to disambiguate ‘*t*’ from ‘*g*’, a finding that mirrored those made for human listeners. The letter ‘*t*’ is also more frequently mislabeled when the AVSR has access to the synthetic videos than when no visual signal is available (yellow annotations in Figure 8A and B). The green rectangles visible in panels A and B of Figure 8 highlight that the synthetic visual representation for the keywords ‘*n*’, ‘*m*’ and ‘*o*’ were a source of confusion for the AVSR, much like for humans.

**Figure 8:**
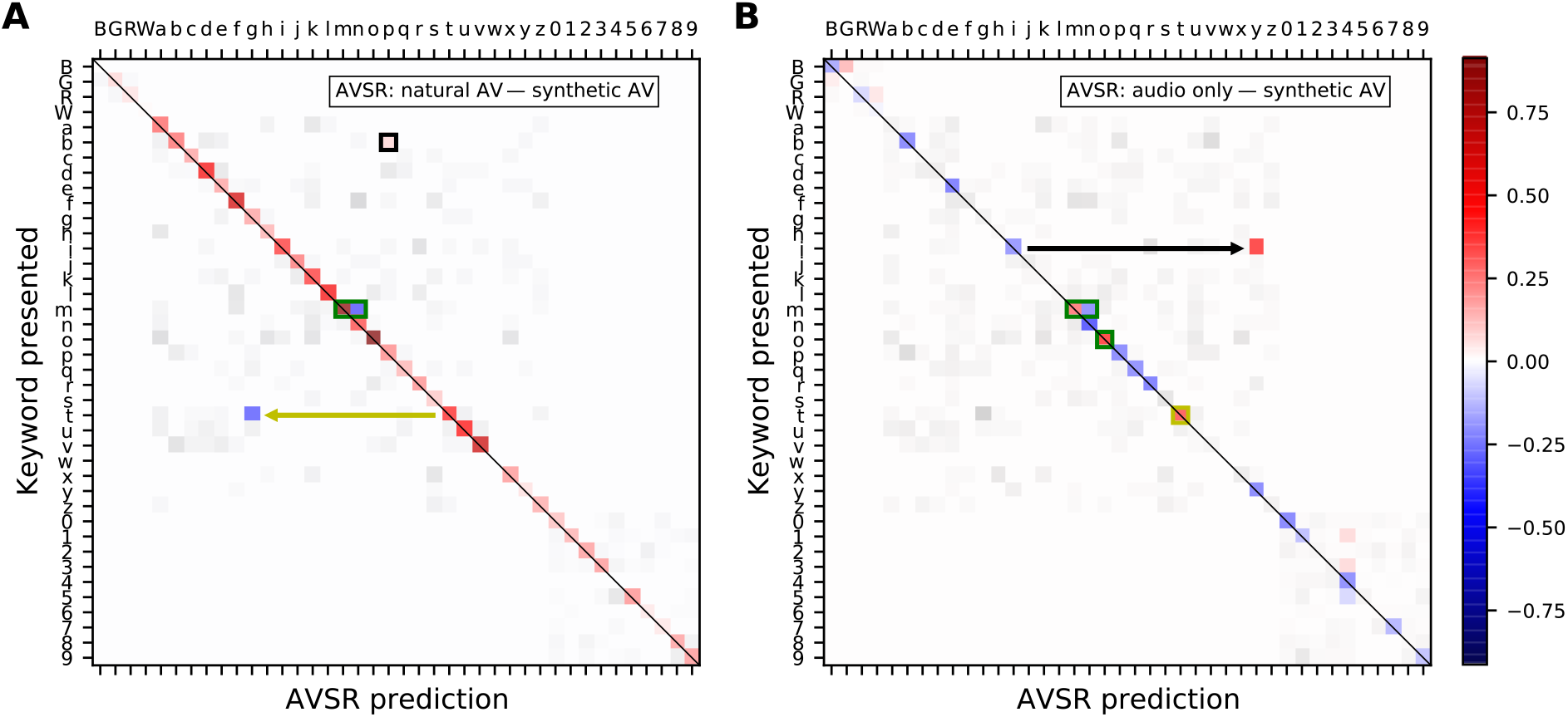
Confusion matrices showing the difference in frequency of presented keyword and participant response pairs between A) the natural audiovisual condition and the synthesized audiovisual condition and B) the audio-only condition and the synthesized audiovisual condition, in the AVSR predictions. The color keywords are represented by their first letter in uppercase, those corresponding to a letter are shown in lowercase, and the number keywords by their digit. The magnitude of statistically significant pairs is highlighted in color (bootstrapped permutation test, alpha=0.05, FDR corrected). In particular, a blue element indicates that the AVSR was significantly more likely to yield the corresponding prediction as a result of having access to a synthesized visual signal as opposed to A) a natural video and B) no video.

Nonetheless, the black annotations in Figure 8 highlight that the synthetic videos had a significantly lower chance to induce the AVSR to label a ‘*b*’ as a ‘*p*’ than their natural counterparts, and that they significantly decreased the chance that the AVSR labelled ‘*i*’ as ‘*y*’ when compared to the audio-only condition.

## 4 Discussion

To the best of our knowledge, our results provide the first demonstration that end-to-end synthetic facial animations can improve speech-in-noise comprehension in humans. Our finding therefore suggest that facial animations generated from deep neural networks can be employed to aid with communication in noisy environments. A next step towards such a practical application will be to investigate the benefit of the facial animations in people with hearing impairment, such as patients with mild-to-moderate sensorineural hearing loss as well as patients with cochlear implants.

However, our results also showed that the speech-in-noise comprehension is yet higher when listeners see the natural videos. This result contrasts with our other finding that humans cannot distinguish between the real and the synthesized videos, neither when explicitly instructed to do so in an online Turing test nor as a spontaneous judgement while carefully and procedurally attending to the videos in a speech-in-noise task using short sentences. We note, however, that the standardized nature of the sentences in the GRID corpus might have hindered the differentiation between the natural and synthetic videos. On the other hand, the Turing test also employed audiovisual material from the TIMIT and CREMA datasets that offer more realistic speech content, such that the standardized nature of the GRID corpus alone cannot explain the observed lack of differentiation in the Turing test. It therefore appears that the synthetic videos lack certain aspects of the speech information, although the lack of this information is not obvious to human observers.

One clue as to why that may be lies in the choice of discriminators employed in the synthesizer GAN architecture: the GAN was optimized for realism and audiovisual synchrony rather than for speech comprehension. Certain keywords pronounced in combination with alveolar and bilabial nasal consonants such as ‘*n*’ and ‘*m*’ or others pronounced in combination with (palato)alveolar affricates and plosives such as ‘*g*’, ‘*t*’ and ‘*two*’ were poorly disambiguated by the synthetic videos. This finding suggests that the GAN may have avoided the issue of synthesizing labial and coronal visemes featuring complex interactions of tongue, teeth, and lip movements to some extent, for the sake of realism and at the expense of comprehension. Still, the result that these videos disambiguated consonants pronounced in combination with the phoneme /i:/ (letter keywords ‘*b*’ and ‘*p*’) and vowels pronounced in combination with the diphthong /aI/ (letter keywords ‘*i*’ and ‘*y*’) signifies that their effectiveness at improving speech comprehension cannot be due to temporal cues alone but must include categorical cues.

From a different perspective, the synthesized audiovisual signals may aid speech comprehension in two ways. First, access to the visual signal may improve the availability of information to human listeners, allowing the brain to perform internal denoising through multimodal integration. This may be aided by the fact that the visual signals were synthesized from clean speech signals without background noise. Second, the synthesizer may be increasing the signal-to-noise ratio externally by adding information regarding the dynamics of visual speech. Such information would be learned by the GAN during training and can be beneficial in speech-in-noise tasks. The latter conclusion is supported by the results presented by Hegde et al. (2021), who recently showed that hallucinating a visual stream by generating it from the audio input can aid to reduce background noise and increase speech intelligibility. Importantly, they also showed that humans scores on subjective scales such as *quality* and *intelligibility* were higher for speech denoised in such a way. Moreover, our finding that an AVSR performs better when it has access to a synthetic facial motion than when it relies on the speech signal alone also suggests that our synthesized facial animations contain useful speech information. We caution, however, that there exist many unknowns regarding the interaction of the AVSR and the GAN-generated stimuli, limiting the further interpretation of the ASVR’s performance on these stimuli.

As a limitation of our experiment, we did not investigate the effects of different temporal lags between the auditory and the visual signals. In a realistic audiovisual hearing aid scenario, the synthetic video signal would be delayed with respect to the audio, due to the sampling and processing time required. Because the auditory signal is often slightly delayed with respect to the visual signal, this inverse temporal latency could influence the AV benefit. Moreover, we did not investigate the effects of different levels and types of background noise on the ability of the synthesizer to accurately reproduce visual speech. In addition, the highly standardized sentences of the GRID corpus, in which the different keywords occurred at the same timing, meant that dynamic prediction was not required for their comprehension. Our study could therefore not assess the influence of the synthetic facial animations on this important aspect of natural speech-in-noise comprehension.

Therefore, a natural progression of this work will be to perform on-line experiments with noise-hardened versions of the synthesizer, such as that proposed by Eskimez et al. (2020). Further studies will also look at improving the synthesizer model through the implementation of targeted loss models, informed by the findings of the confusion matrix analysis presented here.

Taken together, our results suggest that training a GAN-based model in a self-supervised manner and without the use of phonetic annotations is an effective method to capture the lip dynamics relevant to human audiovisual speech perception in noise. This research paves the way for further understanding of the way speech is processed by humans and for applications in devices such as audiovisual hearing aids.

## 5 Funding Acknowledgements

This research was supported by the Royal British Legion Centre for Blast Injury Studies, by EPSRC grants EP/M026728/1 and EP/R032602/1, as well as by the U.S. Army through project 71931-LS-INT.

## 6 Conflict of Interest

The authors declare that the research was conducted in the absence of any commercial or financial relationships that could be construed as a potential conflict of interest.

## Footnotes

1 Pretrained model available at “https://github.com/DinoMan/speech-driven-animation”

2 The Turing test was made available online at “https://forms.gle/vjFzS4QDU9UzFjDJ9”

## Notes

### Competing Interest Statement

The authors have declared no competing interest.

## References

Sumby, W. N., and Pollack, I. (1954). Visual Contribution to Speech Intelligibility in Noise. The Journal of the Acoustical Society of America 26, 212–215. doi: 10.1121/1.1907309

Ross, L. A., Saint-Amour, D., Leavitt, V. M., Javitt, D. C., and Foxe, J. J. (2007). Do You See What I Am Saying? Exploring Visual Enhancement of Speech Comprehension in Noisy Environments. Cerebral Cortex 17:5, 1147–1153. doi: 10.1093/cercor/bhl024

Puschmann, S., Daeglau, M., Stropahl, M., Mirkovic, B., Rosemann, S., Thiel, C. M., et al (2019). Hearing-impaired listeners show increased audiovisual benefit when listening to speech in noise. NeuroImage 196, 261–268. doi: 10.1016/j.neuroimage.2019.04.017

Crosse, M. J., Di Liberto, G. M., and Lalor, E. C. (2016). Eye can hear clearly now: inverse effectiveness in natural audiovisual speech processing relies on long-term crossmodal temporal integration. J. Neurosci. 36, 9888–9895. doi: 10.1523/JNEUROSCI.1396-16.2016

Stevenson, R.A. and James, T.W. (2009). Audiovisual integration in human superior temporal sulcus: Inverse effectiveness and the neural processing of speech and object recognition. Neuroimage 44(3):1210–23. doi: 10.1016/j.neuroimage.2008.09.034

Meredith, M.A. and Stein, B.E. (1986). Visual, auditory, and somatosensory convergence on cells in superior colliculus results in multisensory integration. Journal of Neurophysiology 56(3):640–62. doi: 10.1152/jn.1986.56.3.640

Chandrasekaran, C., Trubanova, A., Stillittano, S., Caplier, A., and Ghazanfar, A. A. (2009) The Natural Statistics of Audiovisual Speech. PLOS Computational Biology 5(7): e1000436. doi: 10.1371/journal.pcbi.1000436

O’Sullivan, A. E., Crosse, M. J., Di Liberto, G. M., de Cheveigné, A., and Lalor, E. C. (2021). Neurophysiological indices of audiovisual speech processing reveal a hierarchy of multisensory integration effects. Journal of Neuroscience 41 (23), 4991–5003. doi: 10.1523/JNEUROSCI.0906-20.2021

Munhall, K.G., Jones, J. A., Callan, D. E., Kuratate, T., and Vatikiotis-Bateson, E. (2004). Visual Prosody and Speech Intelligibility: Head Movement Improves Auditory Speech Perception. Psychological Science 15,2: 133–37. doi: 10.1111/j.0963-7214.2004.01502010.x

Peelle, J. E. and Sommers, M. S. (2015). Prediction and constraint in audiovisual speech perception. Cortex; a journal devoted to the study of the nervous system and behavior, 68, 169–181. doi: 10.1016/j.cortex.2015.03.006

Hickok, G. and Poeppel, D. (2007). The cortical organization of speech processing. Nat Rev Neurosci 8, 393–402. doi: 10.1038/nrn2113

Kayser, C., Petkov, C. I., Remedios, R., and Logothetis, N. K. (2012). Multisensory Influences on Auditory Processing: Perspectives from fMRI and Electrophysiology. Boca Raton (FL): CRC Press/Taylor & Francis

Schroeder, C. E., Lakatos, P., Kajikawa, Y., Partan, S., and Puce, A. (2008). Neuronal oscillations and visual amplification of speech. Trends in cognitive sciences 12(3) 106–13. doi: 10.1016/j.tics.2008.01.002

Kayser, C., Petkov, C. I., Augath, M., and Logothetis, N. K. (2007). Functional imaging reveals visual modulation of specific fields in auditory cortex. Journal of Neuroscience 27(8) 1824–35. doi: 10.1523/JNEUROSCI.4737-06.2007

O’Sullivan, A. E., Crosse, M. J., Di Liberto, G. M., and. Lalor, E.C. (2017). Visual Cortical Entrainment to Motion and Categorical Speech Features during Silent Lipreading. Frontiers in Human Neuroscience 10:679. doi: 10.3389/fnhum.2016.00679

Kuratate, T., Yehia, H., and Vatikiotis-Bateson, E. (1998). Kinematics-based synthesis of realistic talking faces. In Burnham, D., Robert-Ribes, J., Vatikiotis-Bateson, E. (Eds.), International Conference on Auditory-Visual Speech Processing (AVSP’98): 185–190. Terrigal-Sydney, Australia: Causal Productions

Cohen, M.M. and Massaro, D.W. (1990). Synthesis of visible speech. Behavior Research Methods, Instruments, & Computers 22, 260–263. doi: 10.3758/BF03203157

Le Goff B., Guiard-Marigny T., and Benoît C. (1997) Analysis-Synthesis and Intelligibility of a Talking Face. In: van Santen J.P.H., Olive J.P., Sproat R.W., Hirschberg J. (eds) Progress in Speech Synthesis. Springer, New York, NY. doi: 10.1007/978-1-4612-1894-4_18

Massaro, D. W. and Cohen, M. M. (1990). Perception of Synthesized Audible and Visible Speech. Psychological Science, I, 55–63. doi: 10.1111/j.1467-9280.1990.tb00068.x

Bailly, G., Bérar, M., Elisei, F., and Odisio, M. Audiovisual Speech Synthesis. International Journal of Speech Technology 6, 331–346 (2003). doi: 10.1023/A:1025700715107

Fagel, S. and Sendlmeier, W. (2003). An expandable web-based audiovisual text-to-speech synthesis system. Conference: 8th European Conference on Speech Communication and Technology. EUROSPEECH 2003 -INTERSPEECH 2003, Geneva, Switzerland

Fagel, S. (2004). Video-realistic synthetic speech with a parametric visual speech synthesizer. Conference: 8th International Conference on Spoken Language Processing. INTERSPEECH 2004, Jeju Island, Korea. doi: 10.21437/Interspeech.2004-422

Aller, S. and Meister, E. (2016). Perception of Audiovisual Speech Produced by Human and Virtual Speaker. Human Language Technologies – The Baltic Perspective, Ebook Volume 289, Pages 31–38. doi: 10.3233/978-1-61499-701-6-31

Lidestam, B. and Beskow, J. (2006). Visual Phonemic Ambiguity and Speechreading. Journal of speech, language, and hearing research: JSLHR. 49. 835–47. doi: 10.1044/1092-4388(2006/059)

Agelfors E., Beskow J., Karlsson I., Kewley J., Salvi G., and Thomas N. (2006) User Evaluation of the SYNFACE Talking Head Telephone. In: Miesenberger K., Klaus J., Zagler W.L., Karshmer A.I. (eds) Computers Helping People with Special Needs. ICCHP 2006. Lecture Notes in Computer Science, vol 4061. Springer, Berlin, Heidelberg. doi: 10.1007/11788713_86

Beskow, J., Granstrm, B., and Spens, K. (2002). Articulation Strength - Readability Experiments With A Synthetic Talking Face. TMH-QPSR Vol. 44 - Fonetik 2002

Chung, J. S., Jamaludin, A., and Zisserman, A. (2017). You said that?. British Machine Vision Conference 2017 [Preprint], London, UK. 1705.02966v2

Chen, L., Maddox, R. K., Duan, Z., and Xu, C. (2019). Hierarchical Cross-Modal Talking Face Generation with Dynamic Pixel-Wise Loss. CVPR 2019, Long Beach (CA), USA. doi: 10.1109/CVPR.2019.00802

Vougioukas, K., Petridis, S., and Pantic, M. (2020). Realistic Speech-Driven Facial Animation with GANs. Interantional Journal of Computer Vision 128, 1398–1413. doi: 10.1007/s11263-019-01251-8

Cooke, M., Barker, J., Cunningham, S., and Shao, X. (2006). An audio-visual corpus for speech perception and automatic speech recognition. The Journal of the Acoustical Society of America 120, 2421–2424. doi: 10.1121/1.2229005

Assael, Y. M., Shillingford, B., Whiteson, S., and de Freitas, N. (2016). LipNet: End-to-end sentence-level Lipreading. arXiv [Preprint]. 1611.01599

Garofolo, J. S., Lamel, L. F., Fisher, W. M., Fiscus, J. G., Pallett, D. S., and Dahlgren, N. L. (1993). DARPA TIMIT: (Technical report). National Institute of Standards and Technology. doi:10.6028/nist.ir.4930

Cao, H., Cooper, D. G., Keutmann, M. K., Gur, R. C., Nenkova, A., and Verma, R. (2014). CREMA-D: Crowd-Sourced Emotional Multimodal Actors Dataset. IEEE Transactions on Affective Computing 5 (4:377–390). doi: 10.1109/TAFFC.2014.2336244

Ma, P., Petridis, S., and Pantic, M. (2021). End-to-end audio-visual speech recognition with conformers. International Conference on Acoustics, Speech and Signal Processing 2021 [Preprint], Toronto, Ontario, Canada. 2102.06657v1

Hegde, S. B., Prajwal, K. R., Mukhopadhyay, R., Namboodiri V., and Jawahar, C. V. (2021). Visual Speech Enhancement Without a Real Visual Stream. 2021 IEEE Winter Conference on Applications of Computer Vision, 1925–1934, virtual. doi: 10.1109/WACV48630.2021.00197

Eskimez, S. E., Maddox, R. K., Xu, C., and Duan, Z. (2020). End-To-End Generation of Talking Faces from Noisy Speech. International Conference on Acoustics, Speech and Signal Processing 2020, Barcelona, Spain. doi: 10.1109/ICASSP40776.2020.9054103

